# Sensory neurons expressing the atypical olfactory receptor guanylyl cyclase D are required for the acquisition of odor preferences by mice in diverse social contexts

**DOI:** 10.1101/2020.07.11.198812

**Authors:** Arthur D. Zimmerman, Christina R. Nagy, Steven D. Munger

## Abstract

Animals use social communication to learn important information from conspecifics that can guide appropriate behavioral choices. For example, during the social transmission of food preference (STFP), conspecific semiochemicals detected by mouse olfactory sensory neurons (OSNs) expressing the atypical olfactory receptor guanylyl cyclase D (GC-D+ OSNs) promote the acquisition of food preferences in the recipient animal, mitigating the risk of ingesting food contaminated with toxins or pathogens. However, it is unclear if GC-D+ OSNs mediate preference learning outside this specific context. Here, we report that GC-D+ OSNs are required for the acquisition of odor preferences by both adult and juvenile mice, and that GC-D-dependent preference could be formed for conditionally aversive odors. We used a two-choice olfactory behavioral test to assess odor preferences in adult *Gucy2d* +/+, +/- and -/- mice that encountered novel odors together with GC-D+ OSN stimuli (guanylin family peptides), during social investigation of a live conspecific, or during suckling as pups. *Gucy2d* +/+ and +/-mice (which express functional GC-D), but not *Gucy2d* -/- littermates, successfully acquire a preference for the demonstrated odor in any of these behavioral paradigms. Mice could even acquire a GC-D-dependent preference for odors to which they had recently formed a conditioned aversion. Together, these results demonstrate that GC-D+ OSNs mediate the acquisition of socially-transmitted odor preferences in different social and experiential contexts and at different life stages.

## INTRODUCTION

Intraspecific odor communication guides behaviors as diverse as reproduction, aggression, pathogen avoidance and food choice (Galef 2012; Li and Liberles 2015; Stowers and Kuo 2015). In many cases, an animal’s response to conspecific semiochemicals is largely innate and stereotyped. For example, male silk moths will fly towards a source of the volatile female pheromone bombykol (Hansson 1995). Hermaphrodites of the nematode *C. elegans* produce pheromones that attract males but can promote either aggregation or dispersal of other hermaphrodites, depending on the concentration of the pheromones (Macosko *et al*. 2009). Semiochemicals can also shape more complex behaviors. In mice, chemostimuli present in urine and other excretions can promote male-male aggression (Chamero *et al*. 2007) or serve as a signal of social dominance over subordinates (Thoss *et al*. 2019).

Rats and mice will learn to prefer food with a particular odor after interacting with a known conspecific that ate food containing that odor. Such socially transmitted food preferences (STFPs) (Galef 1985b; Munger *et al*. 2010) typically require social interaction with a live conspecific that has eaten odored food (the demonstrator animal). However, rodents prefer to feed where conspecifics have left urine or fecal deposits (Arakawa *et al*. 2013; Galef and Beck 1985; Galef and Buckley 1996), and STFPs can be acquired when the recipient animal (the observer) smells fecal deposits from an animal that ate the odored food (Arakawa *et al*. 2013). STFPs can be also acquired when a food odor is associated with olfactory stimuli present in the breath (e.g., carbon disulfide, CS_2_) or excreted in urine or feces (e.g., guanylin family peptides) of a demonstrator (Arakawa *et al*. 2013; Bean *et al*. 1988). Our previous studies found that olfactory sensory neurons (OSNs) expressing the atypical olfactory receptor guanylyl cyclase GC-D (GC-D+ OSNs) respond to CS_2_ (Munger *et al*. 2010) and guanylin peptides (Leinders-Zufall *et al*. 2007) with high sensitivity and selectivity, and are required for the acquisition of an STFP in mice (Arakawa *et al*. 2013; Munger *et al*. 2010).

However, while GC-D+ OSNs are clearly required for acquiring an STFP, it was unknown if this specialized part of the mouse main olfactory system mediates other types of odor-dependent preference learning. Using *Gucy2d* gene-targeted mice and several behavioral paradigms, we tested the predictions that GC-D+ OSNs mediate the acquisition of socially transmitted odor preferences; that they are necessary for odor preference learning in diverse behavioral contexts and at different life stages; and that a GC-D+ OSN-dependent odor preference can be established for odors to which the animal is conditionally averse.

## METHODS

### Animals

Mice were cared for in accordance with the NIH Guide for the Care and Use of Laboratory Animals, and all experiments were approved by the University of Florida Institutional Animal Care and Use Committee. *Gucy2d-IRES-Mapt-lacZ* (*Gucy2d*) *+/+, +/-* and *-/-* mice were used in these experiments (Leinders-Zufall *et al*. 2007). *Gucy2d +/+* and *Gucy2d +/-* mice show identical responses in GC-D+ OSNs and in STFP behaviors (Arakawa *et al*. 2013; Kelliher and Munger 2015; Leinders-Zufall *et al*. 2007; Munger *et al*. 2010) and thus were grouped together for analysis. Mice were kept on a 12:12 hour light:dark cycle and housed at 74°F. Mice were group-housed (2-5 mice per cage) with littermates of the same sex and provided *ad libitum* access to food and water when not engaged in behavioral experiments. Measurements of odor preference behaviors were performed on mice between 7-12 weeks of age. All experiments were balanced for sex, and no differences in odor preference behaviors were seen between males and females consistent with earlier studies (Leinders-Zufall *et al*. 2007; Munger *et al*. 2010). All animals were naïve to procedures and only used for a single experiment.

### Odor Preference Testing, Live Demonstrator

*Gucy2d +/+, +/-* and *-/-* mice were pair-housed with same-sex littermates. One mouse was randomly selected to be the “demonstrator” while the other served as the “observer.” Observer mice were habituated 1 hr per day for four consecutive days to placement in a clean, empty cage. The day prior to testing (i.e., day 4 of habituation), observer mice were also acclimated for 15 min to a modified two-port odor apparatus (**Figure 1**)(Jagetia *et al*. 2018). Following habituation, both observer and demonstrator mice were food deprived for 16-20 hrs. On the day of testing, observer mice were moved to a clean cage, while demonstrator mice were given 1 hr in the home cage to feed on crushed rodent chow flavored with 2% cocoa (Hershey’s, Hershey, PA) or 1% cinnamon (McCormick, Hunt Valley, MD). Following feeding, demonstrator mice were placed into the clean cage with the observer mouse, where the mice were given 1 hr to interact.

**Figure 1.**
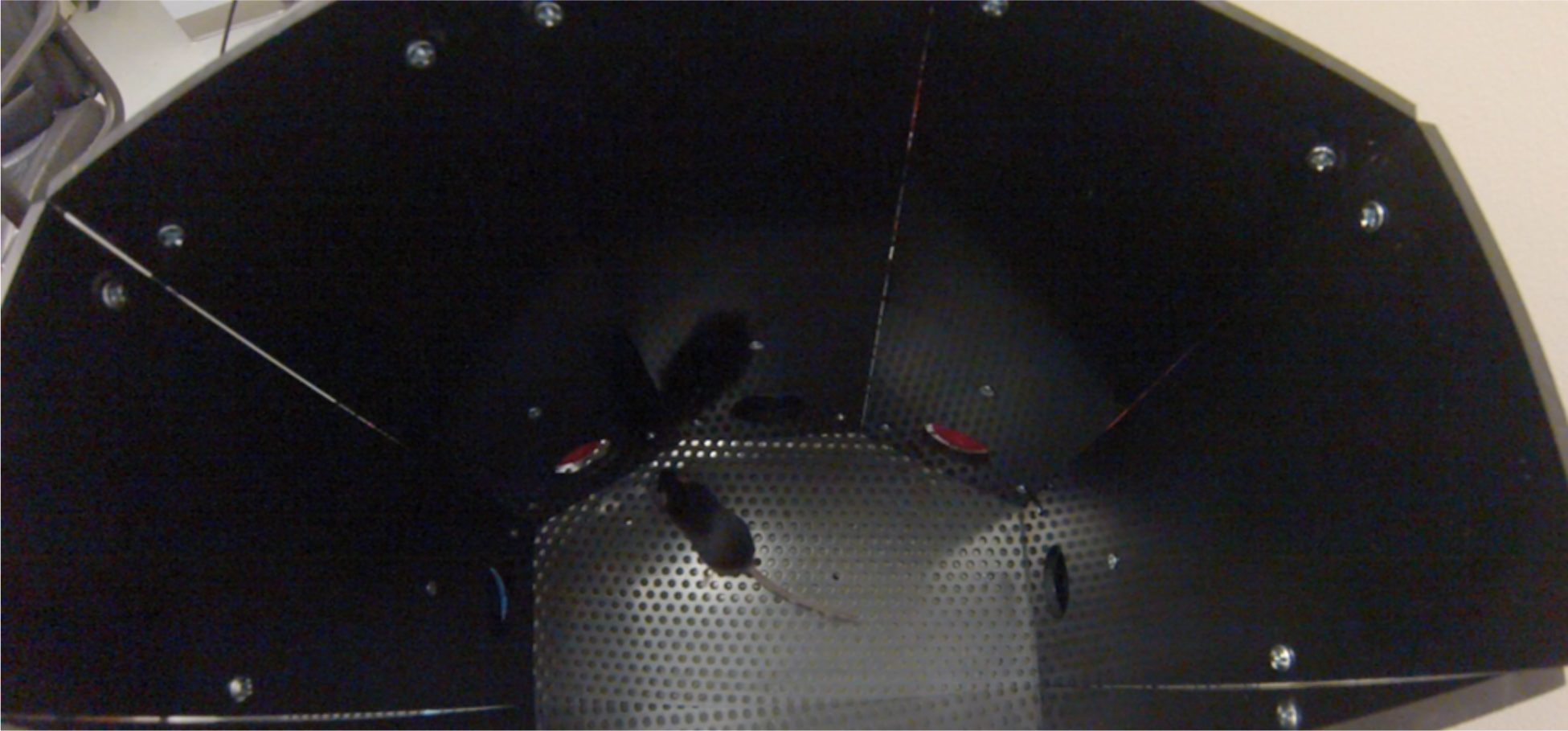
Odor preference testing apparatus. This apparatus, modified from (Jagetia *et al*. 2018), contains two accessible odor ports (red). To sample an odor, a mouse must insert its nose into an odor port and through an infrared beam.

Two hours after interaction with the demonstrator mouse, the observer mouse was placed into the odor preference testing apparatus. This instrument is composed of an open arena with two open odor ports. When mice access a port, they break an infrared beam, resulting in the number and duration of each nose poke being recorded using an Arduino Mega 2560 board on the Arduino Desktop IDE (Integrated Development Environment) software. Odors are drawn from the base of a column just outside the port by a small fan placed at the top of the column, ensuring that odors do not enter the arena. One port contains the demonstrated odor and the other a novel odor (cocoa or cinnamon, counterbalanced across experiments). The mouse is given up to 30 min to explore the arena and the odor ports. A preference ratio was then determined by dividing the amount of time spent sampling the demonstrated odor by the total amount of time spent sampling both odors.

### Odor Preference Testing with Guanylin Peptides

*Gucy2d +/+, +/-* and *-/-* mice were habituated and food deprived similarly as above, but habituation cages also included an empty 60 mm x 15 mm petri dish. On the day of testing, mice were placed in a clean cage for 1 hr where they could interact with a petri dish containing 1 ml of saline with food flavoring (2% cocoa or 1% cinnamon, w/v) in the presence or absence of either guanylin or uroguanylin (50 nM) (Bachem, Torrance, CA; Catalog #H-1342 and H-4148). Two hours following exposure, mice were given a 30 min odor preference test where they could choose to sample either the demonstrated or novel odor (cocoa or cinnamon, counterbalanced across experiments). Experimental measurements and preference ratios were determined as described above.

*Food Preference Testing: Gucy2d +/+, +/-* and *-/-* mice were tested to determine if a GC-D-dependent odor preference could later be manifest as a GC-D-dependent food preference. Briefly, mice were habituated 1 hr per day for four consecutive days to being placed in a clean cage containing an empty 60 mm x 15 mm petri dish. Following habituation on day 4, mice were acclimated for 15 min to the odor preference apparatus the day prior to testing. During the 4-day habituation period, mice were given a diet of only crushed rodent chow to acclimate them for the later food preference test. Following the final day of habituation, mice were food deprived for 16-20 hrs before being placed in a clean cage and given a 1 hr exposure to a saline droplet in a petri dish containing a food flavoring (2% cocoa or 1% cinnamon) in the presence or absence of 50 nM uroguanylin. Mice were then given a 30 min odor preference test as described above. Following odor preference testing, mice were allowed a few hours to eat crushed rodent chow but were then deprived overnight for 23.5 hrs before a food preference test. During the food preference test (Arakawa *et al*. 2013; Beauchamp *et al*. 1983; Kelliher and Munger 2015; Munger *et al*. 2010; Munger *et al*. 2009; Posadas-Andrews and Roper 1983; Ross and Eichenbaum 2006), mice were given a choice between two foods, one containing the demonstrated food flavoring from the previous day and one containing the same novel food flavoring from the odor preference test. A food preference ratio was determined by dividing the amount of food consumed containing the demonstrated food flavoring by the total amount of food consumed.

### Odor Preference Testing After Neonatal Exposure to Maternal Odor

These methods were partially adapted from a study by Fillion and Blass (Fillion and Blass 1986). *Gucy2d +/-* male and female mice were mated to give litters of mixed *Gucy2d* genotype. After birth of the litter, the ventrum of each dam was swabbed once daily with an odor (either 1% citral or 1% eugenol, days P0-P21), and the litters allowed to interact normally with the dam. Upon weaning, pups were separated by sex and group-housed for 4 wks before odor preference testing. At that point, mice were given a 30 min odor preference test where the odor choice was between the demonstrated odor (1% citral or 1% eugenol, present on the dam during suckling) or a novel odor (1% citral or 1% eugenol). Experimental measurements were recorded as above, and preference ratios determined.

### Odor Preference Testing After Conditioned Odor Aversion

These methods were adapted from previously developed procedures for establishing a conditioned odor aversion (Chapuis *et al*. 2009; Slotnick *et al*. 1997). *Gucy2d +/+*, +/- and -/- mice were conditioned to an odor (1% amyl acetate or 1% citral, v/v) via a retronasal procedure. Mice were weighed daily, singly housed and water restricted for 23 hrs throughout the experiment. Mice were transferred to the behavior room for experiments and then returned to the housing room daily. Mice were given 2 ml water for 1 hr for four consecutive days. On the fifth day, mice were given 2 ml water containing amyl acetate or citral (1% v/v). After 5 min, the unconditioned stimulus (either 0.15 M LiCl (Sigma-Aldrich, St. Louis, MO; catalog #203637) or 0.15 M NaCl) was administered via i.p. injection at 6 mEq/kg body weight. Mice were given 2 ml water on day 6, and then conditioned a second time on day 7. The next day, mice were acclimated for 5 min in the odor preference apparatus, and then given 2 ml water. Odor preference testing occurred on day 9. Experimental measures were recorded and preference ratios determined as described above, and mice were given 2 ml of water at the conclusion of testing. On day 10, mice were given 2 ml water for 30 min, and were then moved to another cage where the conditioned odor was presented in saline with uroguanylin (50 nM) for 1 hr. Mice were then fasted for 16–20 hrs. On the final day (day 11), the mice were again given an odor preference test. Odors were counterbalanced across animals, and results were grouped by genotype for statistical analysis (Z scores to assess a significant preference, and 1-way ANOVA to compare genotypes).

### Statistics

The odor preference ratio (PR) was quantified by computing the ratio of time spent sampling the demonstrated odor / the time spent sampling both the demonstrated and novel odors. The food PR was quantified by computing the ratio of the amount of food consumed containing the demonstrated odor / the total amount of food consumed by the observer. Z tests were performed to determine if there was a statistically significant preference for the demonstrated odor or food (a PR of 0.5 indicates no preference), where z = (mean observed odor or food PR – 0.50) / standard error of the mean. Significance between genotypes or experimental conditions was determined by two-way ANOVA and Tukey’s *post-hoc* multiple comparisons test.

## RESULTS

### GC-D-dependent acquisition of odor preferences

We first asked if mice can acquire a GC-D-dependent odor preference from a live conspecific. Observer mice (*Gucy2d +/+, +/-* or *-/-*) were allowed to interact for 1 hr with a littermate (*Gucy2d +/+* or *+/-*) that had just consumed rodent chow containing an added food flavoring (either 2% cocoa or 1% cinnamon). Two hours after the end of this interaction period, observer mice were placed in a two-port nose-poke apparatus (**Figure 1**) to assess the animals’ preference for the demonstrated odor or a novel odor. *Gucy2d +/+* and *+/-* mice, but not *Gucy2d* -/- mice, exhibited a significant preference for the demonstrated odor (p<0.05; **Figure 2**) regardless of which was the demonstrated odor (p=0.97). *Gucy2d +/+* and *+/-* mice spent more total time investigating the demonstrated odor (investigation time per port) and initiated more sampling bouts (i.e., nose pokes) for the demonstrated odor than did *Gucy2d -/-* mice (**Table 1**). There was no difference across genotypes in either the investigation time per sampling bout (i.e., time per nose poke) or the total investigation time across the entire trial (**Table 1**). These data indicate that mice can acquire an odor preference from conspecifics and that this preference acquisition is GC-D-dependent.

**Table 1.**
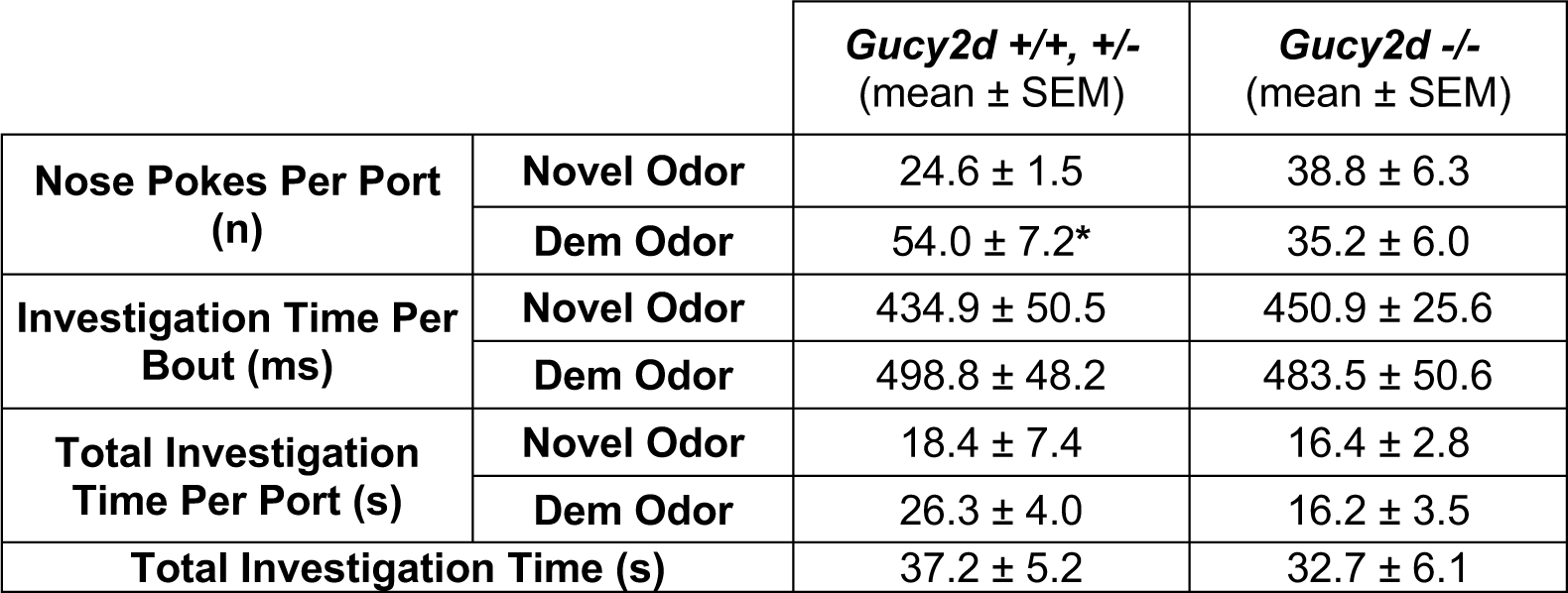
Odor investigation by mice exposed to a live demonstrator.

| | | *Gucy2d* *+/+*, *+/-*<br>(mean ± SEM) | *Gucy2d* *-/-*<br>(mean ± SEM) |
| --- | --- | --- | --- |
| **Nose Pokes Per Port (n)** | **Novel Odor** | 24.6 ± 1.5 | 38.8 ± 6.3 |
| | **Dem Odor** | 54.0 ± 7.2* | 35.2 ± 6.0 |
| **Investigation Time Per Bout (ms)** | **Novel Odor** | 434.9 ± 50.5 | 450.9 ± 25.6 |
| | **Dem Odor** | 498.8 ± 48.2 | 483.5 ± 50.6 |
| **Total Investigation Time Per Port (s)** | **Novel Odor** | 18.4 ± 7.4 | 16.4 ± 2.8 |
| | **Dem Odor** | 26.3 ± 4.0 | 16.2 ± 3.5 |
| **Total Investigation Time (s)** | | 37.2 ± 5.2 | 32.7 ± 6.1 |

**Figure 2.**
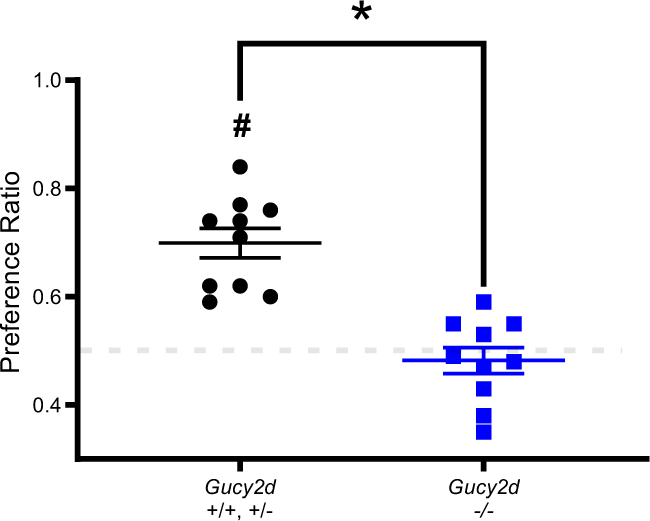
Acquisition of a live demonstrator-dependent socially transmitted odor preference requires GC-D. *Gucy2d +/+* or *+/-* mice (**black circles**), but not *Gucy2d* -/- mice (**blue squares**), exposed to a live demonstrator that had just consumed rodent chow containing an added food odor (2% cocoa or 1% cinnamon) develop an odor preference for the demonstrated odor (n=10 each). *, t-test: p<0.05; #, Z test: p<0.05. Dashed line = no preference.

We next asked if mice can acquire a preference for an odor that is encountered together with olfactory stimuli that specifically activate GC-D+ OSNs. The guanylin-family peptides uroguanylin and guanylin exclusively activate GC-D+ OSNs in the mouse main olfactory epithelium (Leinders-Zufall *et al*. 2007), and their pairing with a general odor promotes the acquisition of a GC-D-dependent food preference (Arakawa *et al*. 2013; Kelliher and Munger 2015). Observer mice were allowed to explore saline droplets that contained a source of general odor (either 2% cocoa or 1% cinnamon, w/v) with or without a guanylin-family peptide (50 nM). *Gucy2d* ^*+/+*^ and ^*+/-*^ mice exposed to a general odor plus either uroguanylin or guanylin exhibited a significant preference for the demonstrated odor (**Figures 3A, B**). Mice failed to acquire an odor preference if they were not exposed to a guanylin peptide or if they did not express GC-D (uroguanylin: p=0.81; guanylin: p>0.99; *Gucy2d* ^-/-^ mice). As with observer mice exposed to live demonstrators, *Gucy2d +/+* and *+/-* mice exposed to the guanylin peptides spent more total time investigating the demonstrated odor and initiated more sampling bouts for the demonstrated odor than did mice that were exposed to only the general odor or did *Gucy2d -/-* mice (**Table 2**). There was also no difference in either the investigation time per sampling bout or the total investigation time across the entire trial across genotypes or peptide exposure groups (**Table 2**). These data indicate that specific activation of GC-D+ OSNs is sufficient to promote the acquisition of a GC-D-dependent odor preference.

**Table 2.** Odor investigation by mice exposed to guanylin peptides.

|  |  | *Gucy2d* +/+, +/-<br>(mean ± SEM) |  | *Gucy2d* -/-<br>(mean ± SEM) |  |
| --- | --- | --- | --- | --- | --- |
|  |  | Odor | Odor +<br>peptide | Odor | Odor +<br>peptide |
| Nose Pokes Per Port (n) | Novel Odor | 44.0 ± 5.6 | 29.7 ± 4.1 | 37.2 ± 7.1 | 32.3 ± 4.3 |
|  | Dem Odor | 45.1 ± 7.3 | 47.7 ± 5.0 | 35.0 ± 7.1 | 26.6 ± 4.9 |
| Investigation Time Per Bout (ms) | Novel Odor | 600.2 ± 48.3 | 507.8 ± 53.0 | 520.5 ± 46.2 | 490.3 ± 35.3 |
|  | Dem Odor | 529.3 ± 55.1 | 550.1 ± 56.3 | 550.0 ± 50.2 | 541.5 ± 37.9 |
| Total Investigation Time Per Port (s) | Novel Odor | 24.6 ± 2.2 | 12.9 ± 1.7 | 17.5 ± 2.2 | 15.5 ± 2.0 |
|  | Dem Odor | 21.1 ± 2.8 | 24.2 ± 2.6* | 16.7 ± 1.7 | 13.6 ± 2.3 |
| Total Investigation Time (s) |  | 45.6 ± 4.5 | 37.2 ± 4.0 | 34.2 ± 3.8 | 29.06 ± 4.1 |

**Figure 3.**
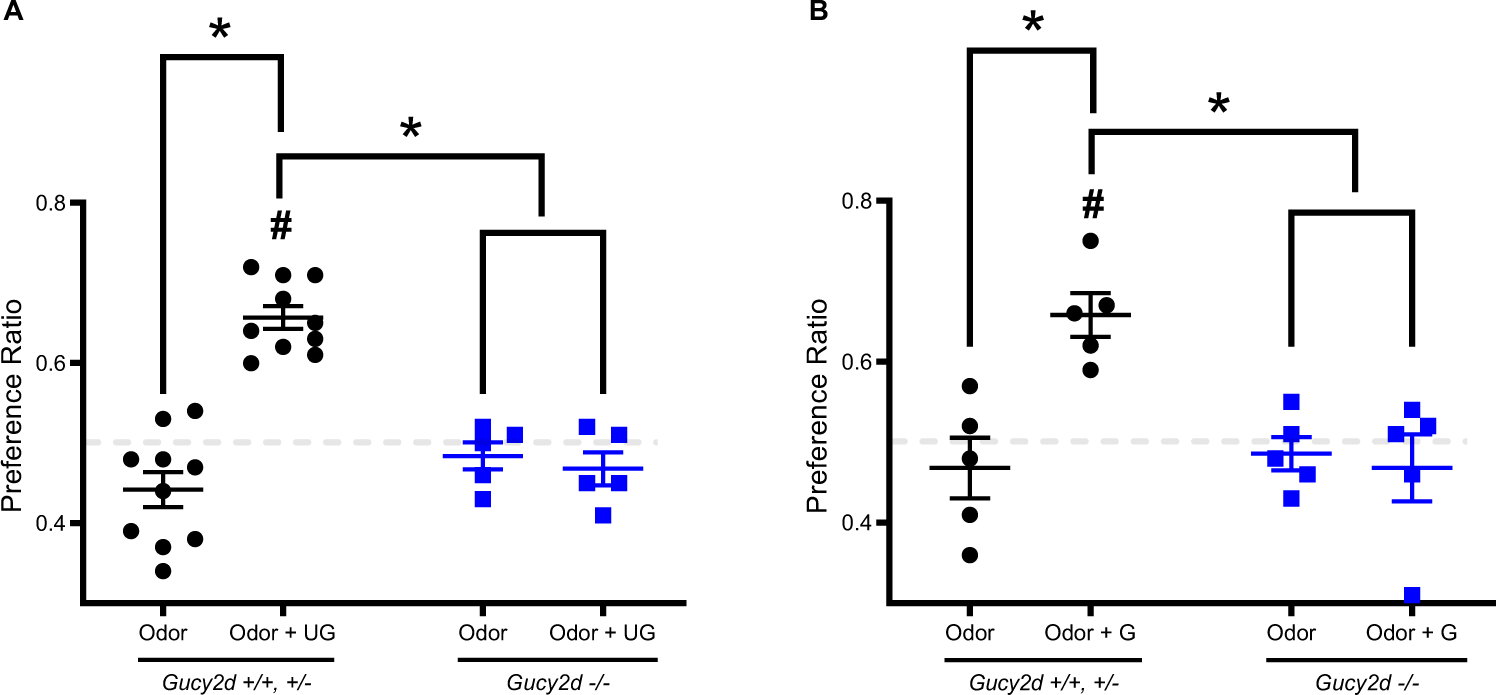
Acquisition of a uroguanylin-or guanylin-dependent socially transmitted odor preference requires GC-D. *Gucy2d +/+* or *+/-* mice (**black circles**), but not *Gucy2d* -/- mice (**blue squares**), exposed to a saline droplet containing an added food odor (2% cocoa or 1% cinnamon) develop an odor preference for the demonstrated odor only in the presence of **(A)** 50 nM uroguanylin (UG) (n=5-10; *: ANOVA: F_1,26_=30.68, p<0.05) or (**B**) guanylin (G) (n=5, *, ANOVA: F_1,16_=10.04, p<0.05). #, Z test: p<0.05. Dashed line = no preference.

### GC-D-dependent odor preferences are associated with an acquired food preference

It is unclear whether mice who have first expressed an odor preference in an odor choice task will still display this preference in a different behavioral context. To address this issue, we performed a food preference test in mice after they had undergone an odor preference test. Similar to results seen in **Figure 2**, *Gucy2d +/+* and *+/-* mice, but not -/- mice, displayed a preference for the demonstrated odor (**Figure 4**). These preferences (or lack of) were maintained the following day when mice were given the choice of foods containing either the demonstrated or a novel odor (**Figure 4**). These data indicate that GC-D-dependent odor preferences can be manifest through distinct behaviors.

**Figure 4.**
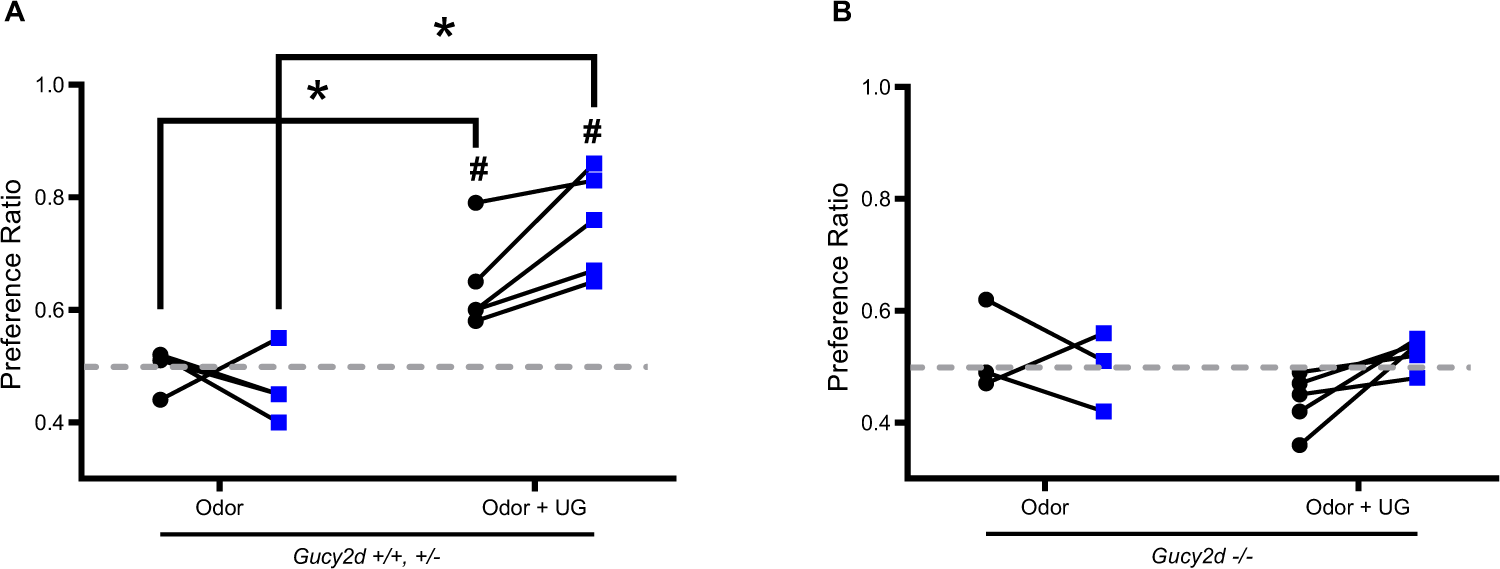
Mice demonstrating UG-dependent odor preferences exhibit equivalent food preferences. *Gucy2d* ^*+/+*^ or ^*+/-*^ mice **(A)**, but not *Gucy2d* ^-/-^ mice **(B)**, exposed to a saline droplet containing an added food odor (2% cocoa or 1% cinnamon) develop an odor preference (**black circles**) for the demonstrated odor only in the presence of 50 nM uroguanylin (UG). *, ANOVA: F_1,14_=13.82 (odor X odor + UG), p<0.05; #, Z test: p<0.05. When given a subsequent food preference test, *Gucy2d* ^*+/+*^ or ^*+/-*^ mice, but not *Gucy2d* ^-/-^ mice, exhibit an equivalent preference for food containing the demonstrated odor (**blue squares**). *, ANOVA: F_1,14_=17.48, p<0.05 (odor X odor + UG). #, Z test: p<0.05. n=3-5. Dashed line = no preference.

### Neonatal GC-D-dependent preferences for odors encountered during maternal interactions are maintained into adulthood

It was unknown whether GC-D-expressing OSNs are important for the acquisition of odor preferences in other behavioral contexts or at other developmental stages. We exposed neonatal mice to monomolecular odors prior to weaning by painting the ventrum of nursing dams with either 1% citral or 1% eugenol (postnatal days 0 - 21). After pups had reached 7 weeks of age, they were then tested for a preference for either citral or eugenol. *Gucy2d +/+* and *+/-* mice, but not -/- mice, showed a preference for the demonstrated odor (p<0.05; **Figure 5**). *Gucy2d +/+* and *+/-* mice showed greater total investigation time, investigation time per bout, and number of nose pokes for the demonstrated odor than for the novel odor (p<0.05; **Table 3**). *Gucy2d* -*/-* mice showed no difference in any of these parameters for the demonstrated or novel odors, and there were no significant differences in total investigation time across genotypes (**Table 3**). Together, these results show that a GC-D-dependent odor preference acquired as a neonate is maintained into adulthood, and indicate that GC-D-expressing OSNs and the olfactory subsystem of which they are a part play a critical role in odor preference learning across multiple social contexts and developmental stages.

**Table 3.** Odor investigation by mice exposed to maternal odors.

|  |  | *Gucy2d*<sup>+/+, +/-</sup><br>(mean ± SEM) | *Gucy2d*<sup>-/-</sup><br>(mean ± SEM) |
| --- | --- | --- | --- |
| **Nose Pokes Per Port (n)** | **Novel Odor** | 22.75 ± 1.47 | 31.09 ± 3.28 |
| **Investigation Time Per Bout (ms)** | **Dem Odor** | 39.19 ± 4.16* | 29.82 ± 2.38 |
| **Investigation Time Per Bout (ms)** | **Novel Odor** | 347.63 ± 19.15 | 400.17 ± 25.26 |
|  | **Dem Odor** | 446.53 ± 23.04* | 394.39 ± 37.60 |
| **Total Investigation Time Per Port (s)** | **Novel Odor** | 7.87 ± 0.64 | 12.02 ± 1.20 |
|  | **Dem Odor** | 17.06 ± 1.82* | 11.52 ± 1.12 |
| **Total Investigation Time (s)** |  | 24.93 ± 2.04 | 23.53 ± 2.22 |

**Figure 5.**
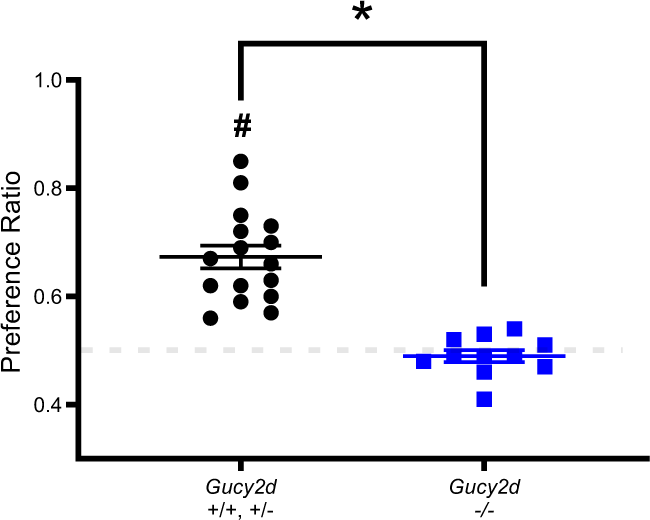
Acquisition of an odor preference following exposure to a maternal odor requires GC-D. *Gucy2d* ^*+/+*^ or ^*+/-*^ mice (**black circles**), but not *Gucy2d* ^-/-^ mice (**blue squares**), exposed to an odor scented on their mother during suckling (1% citral or 1% eugenol) develop an odor preference for the swabbed odor (n=11-16) *, t-test: p<0.05; #, Z test: p<0.05. Dashed line = no preference.

### Preference acquisition for a conditionally aversive odorant

We next asked if mice were able to form a preference for an odorant to which they have been made conditionally averse. Mice were conditioned to avoid an odor (1% amyl acetate or 1% citral; v/v; delivered via water bottle) through association with gastrointestinal malaise (LiCl injection). Mice developed an aversion to either odor when it was paired with a LiCl injection, but not with a control NaCl injection, as seen in an odor preference test (**Figure 6**; p<0.05). Upon re-exposure to the same odor, this time together with uroguanylin, *Gucy2d* +/+ and +/-mice, but not -/- mice, were able to display a preference for that odor (**Figure 6**; p<0.05). These results show that mice can form a GC-D-dependent preference for a conditionally aversive odor.

**Figure 6.**
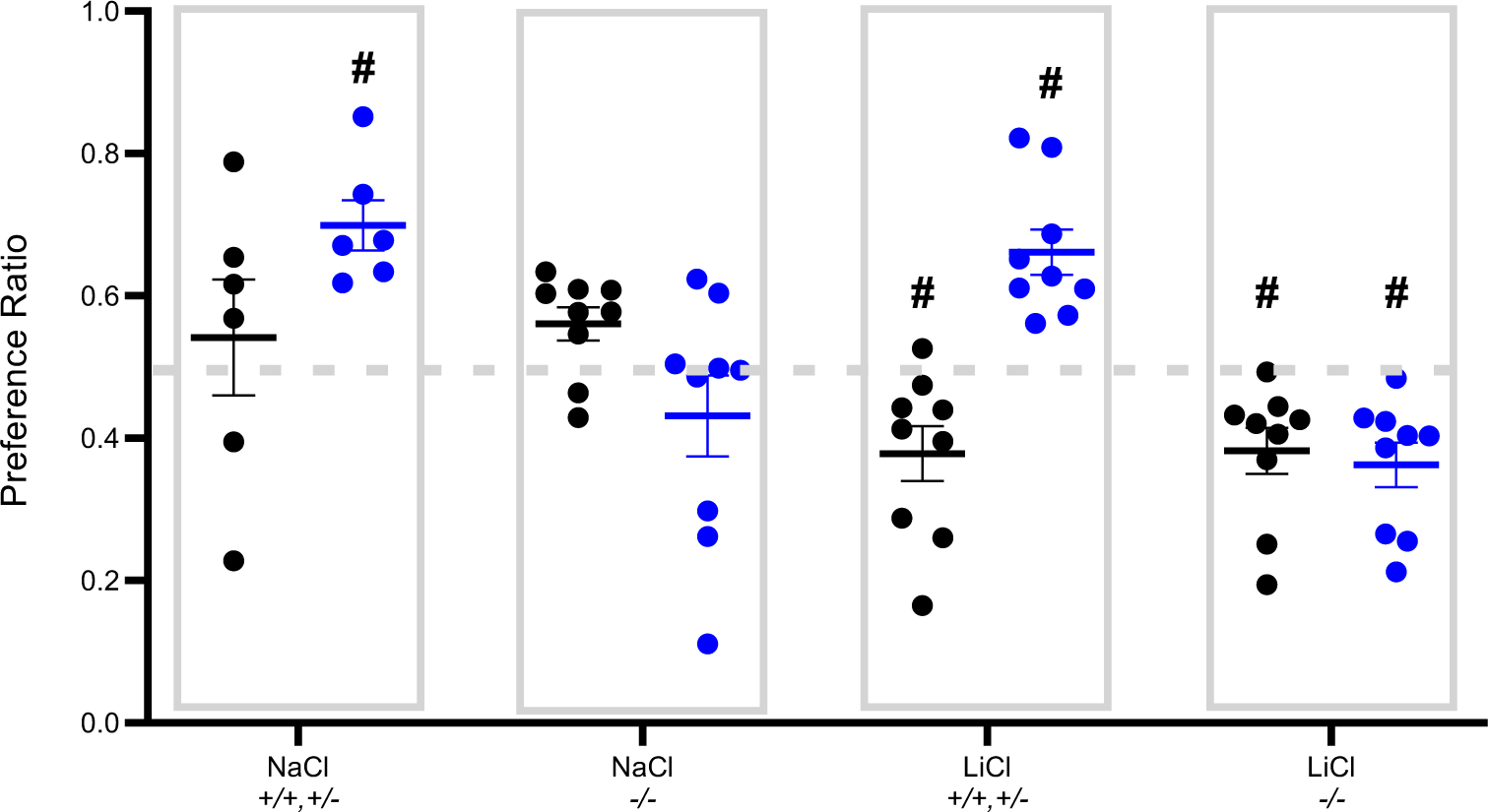
Acquisition of an odor preference following conditioned odor aversion requires GC-D. Injections of LiCl, but not NaCl, paired with an odor (1% citral or 1% eugenol) resulted in a conditioned aversion to that odor (**black circles**). Subsequent pairing of the odor with 50 nM uroguanylin (**blue circles)** resulted in an odor preference in *Gucy2d +/+* or *+/-* mice injected with either NaCl or LiCl, but not in *Gucy2d* -/- mice. (n=6-9). #, Z test: p<0.05. Dashed line = no preference.

## DISCUSSION

Whether an STFP is essentially a socially transmitted odor preference that is expressed as a food choice was unclear. For example, observer rats exposed to a demonstrator rat form a preference for foods containing the demonstrated odor but not for nesting materials or nest boxes containing that same odor (Galef *et al*. 1994), suggesting that social influences on food odors following an STFP do not drive a general preference for the odor in other behavioral contexts. Rather, the interpretation was that learned preferences for a food odor either require food consumption or are restricted to feeding behaviors. By contrast, the same group found that rats also prefer to ingest fluids containing a demonstrated odor following fluid deprivation (Galef *et al*. 1984). Here, we found that, in addition to mediating the acquisition of an STFP, GC-D+ OSNs are essential for the acquisition of a socially transmitted odor preference. This odor preference could be acquired from a live demonstrator or by pairing the demonstrated odor with a GC-D+ OSN-specific olfactory stimulus. Furthermore, while this preference could be expressed by the consumption of a chosen food, it need not be. We note that these mice were food restricted prior to the odor preference test, and may therefore be motivated to find food during the preference testing. While *Gucy2d* ^-/-^ mice showed fewer sampling bouts and less sampling time for the demonstrated odor than did mice with functional GC-D+ OSNs, they did not differ in total sampling time or the total number of individual sampling bouts during odor preference testing. Consistent with previous studies in these mice (Leinders-Zufall *et al*. 2007), we can conclude that differences in odor sampling can be attributed to a change in preference and not in general olfactory ability.

Associative odor learning has a critical impact on survival from early infancy. Human infants begin to use associative odor learning during the prenatal period as infants preferentially orient their heads toward their own mother’s amniotic fluid (Schaal *et al*. 1998). During the postnatal period, infants will begin to mouth when they smell their mother’s odor (Sullivan and Toubas 1998). Rodent pups also rely on odor learning for suckling, huddling and home orientation (Alberts 2007; Cheslock *et al*. 2000; Rosenblatt 1983). Odors of foods consumed by female rats can be encountered by fetuses via amniotic fluid or pups via milk during suckling. The pups will maintain preferences for foods containing those odors, even post weaning (Capretta and Rawls 1974; Galef 1985a; Hepper 1988). Here, we modified an experiment that found that neonatal odor exposure influenced odor-dependent sexual behaviors in adult rats (Fillion and Blass 1986) to ask if GC-D-dependent odor preferences in adults could be acquired during neonatal odor exposure. GC-D+ OSNs coalesce into glomeruli in the necklace region of the caudal main olfactory bulb within the first few postnatal days (Walz *et al*. 2007), so should be capable of responding to maternal chemostimuli (e.g., CS_2_, guanylin peptides) during most of the pre-weaning period. Consistent with studies that reported odor preferences can be maintained for weeks after an initial exposure (Galef 1985a; Hepper 1988), we found that the GC-D-dependent odor preference formed by pups during interactions with their mother were strongly maintained at least four weeks later. These results are also consistent with other studies that have shown rats can develop and maintain an odor preference when exposed to an odor on the ventrum of the dam during the preweaning phase (Galef and Kaner 1980; Galef 1982). Together, our findings that both neonates and adults can form GC-D-dependent odor preferences argues that this form of social learning can occur in multiple behavioral contexts, at different life stages, and is not limited to influencing choices about food consumption.

Animals must be able to overcome maladaptive aversions to sensory stimuli, such as those associated with foods, that are normally safe or beneficial (Galef 1989). For example, while bitter-tasting compounds can signal potential toxicity, humans can learn to prefer bitterness in food and drink (Beauchamp and Mennella 2011; Forestell and Mennella 2007). In the context of an STFP, observer rats will acquire a preference for a food to which they had recently formed a conditioned aversion when they are exposed to demonstrator rats who had eaten that same food (Galef 1986). Similarly, prior exposure of an observer rat to a demonstrator that has eaten a specific food will interfere with the subsequent acquisition of a LiCl-induced aversion to that food (Galef 1989). We previously reported that observer mice will continue to prefer a demonstrated food even when that food contains the rodenticide warfarin (Kelliher and Munger 2015). And rats will prefer a less palatable food over a more palatable choice when they are previously exposed to a demonstrator that consumed the less palatable food (Galef 1986). We found that mice were able to form a GC-D-dependent preference for odors to which they had recently formed a conditioned aversion. Therefore, it appears that odor learning mediated by GC-D+ OSNs can overcome at least some other forms of odor learning. It would be interesting to know if mice could also form a GC-D-dependent preference for innately aversive odors, such as predator kairomones, including those that activate other olfactory tissues such as the Gruenberg ganglion or vomeronasal organ. However, the repeated olfactory sampling typically employed in the behavioral paradigms used here are likely to be a challenge for odors that cause stereotyped defensive or flight behaviors.

## ACKNOWLEDGEMENTS AND FUNDING

This work was supported by the National Institute on Deafness and Communication Disorders with grants R01 DC005633 and T32 DC0159914. Behavioral studies were performed in the University of Florida Center for Smell and Taste Behavioral Core Laboratory. The authors declare no conflicts of interest.

